# Estradiol modulates gut microbiota in female *ob/ob* mice fed a high fat diet

**DOI:** 10.1101/612283

**Authors:** Kalpana D Acharya, Xing Gao, Elizabeth P Bless, Jun Chen, Marc J Tetel

## Abstract

Estrogens protect against diet-induced obesity in women and female rodents. In support of these anorectic effects, lack of estrogens in postmenopausal women is associated with weight gain, increasing their risk for cardiovascular diseases and cancer. Estrogens act with leptin, a satiety hormone encoded by the *ob* gene, to regulate energy homeostasis in females. Leptin-deficient mice (*ob/ob*) exhibit morbid obesity and insulin resistance. In addition to estrogens and leptin, the gut microbiome (gut microbes and their metabolites), is critical in regulating energy metabolism. The present study investigates whether estrogens and leptin modulate gut microbiota in ovariectomized *ob/ob* (obese) or heterozygote (lean) control mice fed a high-fat diet (HFD) that received either 17β-Estradiol (E2) or vehicle implants. E2 attenuated weight gain in both genotypes compared to vehicle counterparts. Moreover, both obesity (*ob/ob* mice) and E2 reduced gut microbial diversity. *ob/ob* mice exhibited lower species richness than control mice, while E2-treated mice had reduced evenness compared to vehicle mice. Regarding taxa, E2 treatment was associated with higher abundances of the family S24-7. Leptin was associated with higher abundances of Coriobacteriaceae, *Clostridium* and *Lactobacillus*. E2 and leptin had overlapping effects on relative abundances of some taxa, suggesting that interaction of these hormones is important in gut microbial homeostasis. Taken together, these findings suggest that E2 and leptin profoundly alter the gut microbiota of HFD-fed female mice. Understanding the function of E2 and leptin in regulating gut microbiota will allow the development of therapies targeting the gut microbiome for hormone-dependent metabolic disorders in women.

## Introduction

Estrogens profoundly influence energy homeostasis [1-3], as well as reproductive physiology and behavior [4-6]. Estrogens reduce food intake, attenuate body weight gain and adiposity, and increase physical activity in both humans and rodents [1, 2]. Postmenopausal women have lower levels of circulating estrogens and an increased tendency to gain fat weight, which increases their risk for obesity, cardiovascular disease, stroke, and type 2 diabetes [7-9]. Similarly, in mice on a high-fat diet (HFD), ovariectomy increases energy intake and the development of obesity, while estradiol (E2) treatment prevents weight gain [2, 10-13], indicating that estrogens protect against HFD-induced obesity.

Leptin is a peptide hormone secreted primarily by adipocytes, which acts in the brain to stimulate metabolism, promote satiety, and regulate fat storage [14-16]. A mutation in the *ob* gene that encodes leptin results in mice lacking the hormone (*ob/ob)* [17]. While phenotypically normal at birth, *ob/ob* mice quickly develop obesity and diabetes [18]. Additionally, *ob/ob* mice exhibit increased food intake and decreased physical activity, energy metabolism, and body temperature compared to lean controls, making *ob/ob* mice an excellent genetic model of obesity [19-21]. Administering leptin to adult *ob/ob* mice reverses these effects by decreasing food intake, increasing energy output and decreasing circulating levels of glucose and insulin [22, 23].

Leptin and estrogen signaling pathways interact to influence reproduction and energy metabolism. High levels of E2 are associated with increased leptin sensitivity in both male and female rodents [24]. In mice, ovariectomy decreases leptin sensitivity, but can be restored by E2 administration [25]. While female *ob/ob* mice do not have an estrous cycle, leptin administration restores fertility, including successful ovulation, pregnancy, and birth, indicating the profound effects of leptin on reproduction [26]. While these findings indicate functional interactions between E2 and leptin, mechanisms by which these hormones interact to regulate energy metabolism are not well understood.

The gut microbiome, which is composed primarily of the bacteria in the intestinal tract and their metabolites, has profound effects on energy metabolism [27]. Changes in body weight have been associated with changes in gut microbiota diversity in humans and rodents [28]. For example, the transfer of gut microbiota from obese human or mice donors results in an obese phenotype in recipient mice [29-31]. The gut microbiome aids in digestion and absorption of macro- and micro-nutrients from food [32]. Gut microbiota are also essential modulators of host immune homeostasis. Protective polysaccharides produced from the break-down of dietary fibers attenuate inflammation [33, 34]. Finally, gut microbiota can synthesize and metabolize neurotransmitters and hormones to alter host physiology [35-37].

A variety of factors, including host genetics, diet, stress, and gonadal hormones can alter the gut microbiome [36-40]. Sex differences in gut microbiota have been reported in humans and rodents [41]. In a European population, higher levels of Bacteroidetes and Prevotella were observed in men compared to women [42]. Male mice exhibited higher abundances of Lachnospiraceae (phylum Firmicutes) and *Parabacteroides* spp. (phylum Bacteroidetes) and Proteobacteria than female mice [36, 43]. Additionally, testosterone administration to female neonatal rats decreased gut microbial diversity during adulthood, and increased the ratio of the two most abundant phyla, Firmicutes and Bacteroidetes [38]. The hormone-dependent changes in gut microbiota were more robust compared to the diet-induced changes in these rats [38]. Ovariectomy also alters gut microbial diversity in adult mice [36, 43, 44]. While obesity and *ob/ob* genotype are associated with a reduction in Bacteroidetes/Firmicutes ratio in humans and male mice, respectively [45-48], the effects of leptin on gut microbiota have not been studied in females. Since estrogens and leptin interact to affect female metabolic homeostasis [10, 25], it is important to understand the effects of leptin and its interaction with estrogens, on gut microbiota in females. Thus, using a mouse model of obesity (*ob/ob*), the present study tested the hypothesis that E2 and leptin alter gut microbiota and energy homeostasis in female mice.

## MATERIALS AND METHODS

### Animals

Seven week-old lean heterozygote (Het) and obese *ob/ob* (leptin-deficient) mice (24 mice/genotype) were purchased from Jackson Laboratory (Bar Harbor, Maine), kept on a 12:12 light:dark cycle and allowed to acclimate at the Wellesley College animal facility for a week. Het mice, which have a metabolic phenotype similar to wildtype mice as evidenced by normal serum glucose and insulin, body temperature, and energy expenditure [19], were used as controls [21] to account for genetic background. While phenotypically very similar, Het mice have lower leptin levels than wildtype mice [49]. All mice were ovariectomized (OVX) and subcutaneously implanted with a silastic capsule containing 17/*β*-Estradiol (E2, #E8875, Sigma; 50µg in 25µl of 5% ethanol/sesame oil) or Veh (5% ethanol/sesame oil) resulting in the following four experimental groups: control E2, 2) control Veh, 3) *ob/ob* E2, and 4) *ob/ob* Veh. Following surgery, mice were pair-housed with a mouse of the same genotype and treatment.

As part of another study to identify newborn cells, on day 7 mice underwent intracranial surgery for the implantation of an intraventricular cannula attached to a subcutaneous osmotic pump filled with bromodeoxyuridine. The osmotic pumps were removed after 10 days. On day 35, animals received an intraperitoneal injection of leptin (5 mg/kg) 45 minutes prior to being euthanized for assessment of acute leptin response in the brain, unrelated to the present study. While possible, it is unlikely that leptin administration 45 minutes prior to sacrifice would have effects on gut microbiota. All procedures were approved by the Institutional Animal Care and Use Committee of Wellesley College.

### Food intake and body weight measurements and fecal sample collection

Following OVX (day 0), all mice were maintained on a standard diet for three days (13.5% calories from fat, Purina, #5001) before switching to a HFD (58.4% calories from fat Harlan Teklad, #03584) on day 4 and maintained on a HFD for the remainder of the study. Throughout the experiment, food intake and body weights were recorded every four days from heterozygote control (Het) E2 (n=13), Het Veh (n=12), *ob/ob* E2 (n=8) and *ob/ob* Veh (n=7) mice. Food intake was calculated per cage, from a total of Het E2 (n=6), Het Veh (n=6), *ob/ob* E2 (n=6) and *ob/ob* Veh (n=6) cages, with each cage containing two mice of the same treatment. For cages in which one mouse died, the amount of food eaten by the remaining mouse was doubled to match with cages containing two mice. Similarly, for microbiota analysis, fecal samples from each cage was counted as n=1, resulting in Het E2 (n=6), Het Veh (n=6), *ob/ob* E2 (n=6) and *ob/ob* Veh (n=6). Fecal samples were collected on days 4, 7, 15, 23, 31, and 35 and immediately stored at −80°C.

### DNA extraction from fecal samples, 16S rDNA sequencing and bioinformatics processing

DNA was extracted from fecal samples using a MoBio PowerSoil® DNA Isolation Kit (Valencia, CA) with minor adjustments to the manufacturer’s protocol. A 5-minute incubation with the elution buffer before centrifugation was added to increase the DNA yield. The quality and quantity of the DNA samples were measured using Nanodrop (Thermo Scientific, Waltham, MA). The samples were stored at −20°C until sequencing.

The V3-V4 region of the 16S rDNA was amplified using the following universal 16S rDNA primers: forward 341F (5’-CCTACGGGAGGCAGCAG-3’) and reverse 806R (5’-GGACTACHVGGGTWTCTAAT-3’) with sequence adapters on both primers and sample-specific Golay barcodes on the reverse primer [50]. The PCR products were quantified by PicoGreen (Invitrogen, Carlsbad, CA) using a plate reader. After quantification, amplicons were pooled in equal concentrations, cleaned up using UltraClean PCR Clean-Up Kit (MoBio, Carlsbad, CA), and again quantified using the Qubit (Invitrogen, Carlsbad, CA). The pooled samples were then sequenced using paired-end v2 chemistry using Illumina Miseq sequencing technology (Illumina, San Diego, CA) at the Microbiome Core, Mayo Clinic (Rochester, Minnesota).

Paired R1 and R2 sequence reads were then processed via the *hybrid-denovo* bioinformatics pipeline, which clustered a mixture of good-quality paired-end and single-end reads into operational taxonomic units (OTUs) at 97% similarity level [51]. OTUs were assigned taxonomy using the RDP classifier trained on the GreenGenes database (v13.5) [52]. A phylogenetic tree based on FastTree algorithm was constructed based on the OTU representative sequences [53]. Singleton OTUs as well as samples with less than 2,000 reads were removed from downstream analysis.

### Statistical analysis

#### Food intake and body weight data analysis

Repeated measures ANOVA (SPSS, v.24) was performed to examine the effects of treatment and genotype on food intake and body weight over time. After a main effect was confirmed, ANOVA without corrections was performed on measures from each day to identify the days with effects of one or both variables. One–way ANOVAs were then conducted to identify differences between specific groups when effects of treatment or genotype were present. Differences were considered statistically significant at p < 0.05.

#### 16S rDNA sequence analysis

Analyses were first performed on the aggregated data, in which sequence reads from each mouse across all days were aggregated. To study specific longitudinal trends, stratified analyses on individual days were also performed if needed.

#### Diversity analysis

Both α-diversity and β-diversity were analyzed on the rarefied OTU data. α-diversity (within-sample diversity) reflects species richness and evenness within the microbial populations. Two representative α-diversity measures were investigated: the observed number of OTUs, an index of the species richness, and the Pielou’s evenness index [54]. A multiple linear regression model (“lm” function in R) was used to test the association between α-diversity (outcome) and treatment/genotype (covariates, both included in the model).

β-diversity (between-sample diversity) reflects the shared diversity between bacterial populations in terms of ecological distances; pair-wise distance measure allows quantification of the overall compositional difference between samples. Different β-diversity measures provide distinctive views of the community structure. The β-diversity measures were calculated using Bray-Curtis Dissimilarity, which measures differences in bacterial composition based on taxa abundances (“vegdist” function in the R “vegan” package, v2.4-3). To test the association between β-diversity measures and treatment or genotype, we used PERMANOVA (999 permutations, “adonis” function in the R “vegan” package, v2.4-3) when adjusting the effect of the other covariate. Ordination plots were generated using principal coordinate analysis (PCoA) on the distance matrix (“cmdscale” function in R) [55].

#### Taxa level analysis

Differential abundance analyses were performed at the phylum, class, order, family and genus levels. Taxa with prevalence less than 10% or with a maximum proportion less than 0.2% were excluded from analysis to reduce the number of tests. An overdispersed Poisson model was fitted to the taxa counts with treatment and genotype as covariates (“glm” function in R) [56]. Wald test was used to assess significance. To account for variable sequencing depths, the GMPR size factor was estimated and used as an offset (log scale) in the regression model [57]. False discovery rate (FDR) control (B-H procedure, ‘p.adjust’ in R) was used to correct for multiple testing at each taxonomical level, and FDR-adjusted p-values (q-values) < 0.1 were considered significant [58]. The differential taxa were visualized on a cladogram using GraPhlAn [59]. All statistical analyses were performed in R (v. 3.3.2, R Development Core Team).

## Results

### Estradiol and ob/ob genotype altered weight gain and food intake in female mice on a HFD

Body weights were collected every four days from the four experimental groups: control E2 (n=13), control Veh (n=12), *ob/ob* E2 (n=8), and *ob/ob* Veh (n=8). Longitudinal analysis of weight change showed that E2 decreased weight gain (F=15.67, p<0.001; ANOVA) in female mice fed a HFD. E2 treatment attenuated weight gain from day 11 through the end of the study (Figures 1A and 1B). Within-genotype comparison showed that E2 attenuated weight gain in Het mice from days 11-35, compared to Veh mice. For *ob/ob* mice, E2 reduced weight gain on days 11, 15 and 27-35, and showed strong trends on days 19 (p=0.059) and 23 (p=0.063). Because the Het and *ob/ob* mice had different weights at the beginning of the study, the % weight gain was also analyzed to remove the bias due to existing differences in weights prior to hormone and diet manipulation. Similar to the effects on body weight, E2 treatment reduced % weight gain from days 11-35. In particular, E2-treated Het mice gained less % weight than Veh Het mice on days 11-35 (Figure 1B). Within the *ob/ob* mice, E2 decreased % weight gain on days 7-15 and 27-35.

**Figure 1.**
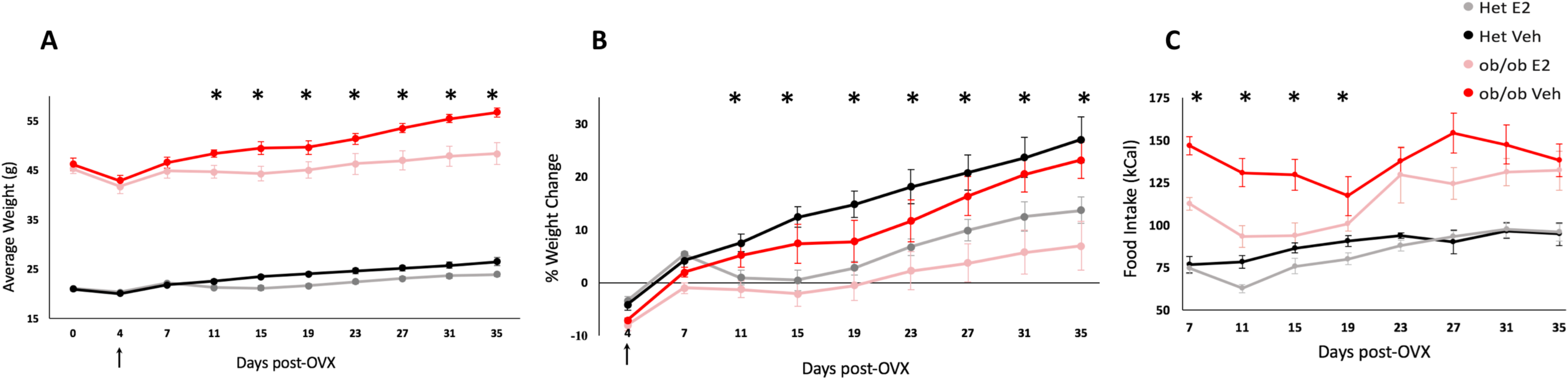
Estradiol and leptin alter weight gain and food intake in adult female *ob/ob* and control mice. (A) Average weight, (B) percent weight gain and (C) food intake (2 mice/cage) of heterozygote controls (Het) E2 (n=13), Het Veh (n=12), *ob/ob* E2 (n=8) and *ob/ob* Veh (n=7) with arrow indicating the start of high fat diet. The effects of genotype on average weight (A) and food intake (C) were present on all days. Days with effects of E2 are denoted by * (p<0.05; ANOVA). Error bars indicate ±SEM.

Genotype also affected weight gain (F=1119.79, p<0.001; ANOVA) and % weight gain on longitudinal measures (F=4.23, p=0.047; ANOVA). *ob/ob* mice weighed more than control Het mice at all time points regardless of E2 treatment. At the start of the study, *ob/ob* mice weighed approximately twice that of the Het mice, which persisted throughout the study. Finally, there was an interaction of E2 and genotype on body weight (F=4.21, p=0.048; ANOVA), with E2-treated Het mice gaining more % weight than E2-treated *ob/ob* mice on days 4 and 7 (Figure 1B). The Veh mice did not differ in % weight gain between the two genotypes.

E2 treatment reduced HFD consumption (F=11.04, p=0.002; ANOVA) from days 7 to 19 (Figure 1C). Analysis within each genotype showed that E2 attenuated food intake in *ob/ob* mice, but not in Hets, on days 7, 11 and 15. Comparison between lean Hets and *ob/ob* mice showed that obesity profoundly increased food intake (F=110.5, p<0.001, ANOVA). Het mice consumed less calories than *ob/ob* mice throughout the study. Within the E2-treated groups, an increase in food intake in *ob/ob* groups compared to E2 Het mice was observed only at the beginning (days 7 and 11) and end (days 27, 31 and 35) of the study. There was an interaction between genotype and treatment on food intake (F=5.52, p=0.029; ANOVA). We also observed a decrease in food intake on day 19 in *ob/ob* Veh mice, but not other groups, following surgery on day 16 for removal of BrdU osmotic pumps.

### Estradiol and ob/ob genotype alter gut microbial diversity during HFD

To assess the impact of E2 treatment and obesity on α-diversity of the gut microbiota, 16S rDNA from fecal samples from *ob/ob* and Het mice were analyzed. The data set contained 137 samples after removing samples with less than 2,000 reads. The 16S rDNA targeted sequencing yielded 25399 reads/sample on average (range 11,154 - 82,733). Clustering of these 16S sequence tags produced 473 OTUs at 97% similarity level. The identified OTUs belonged to 10 phyla, 42 families and 62 genera based on the Greengenes database. During HFD, *ob/ob* mice had a lower number of total identified OTUs than Het mice, indicating that *ob/ob* mice had lower species richness than control mice (Figure 2A; p=0.002, linear regression). In addition, E2 treatment was associated with lower species evenness in both genotypes, suggesting that E2-treated mice have a more heterogeneous distribution of gut microbial communities than Veh mice (Figure 2B; p=0.0008).

**Figure 2.**
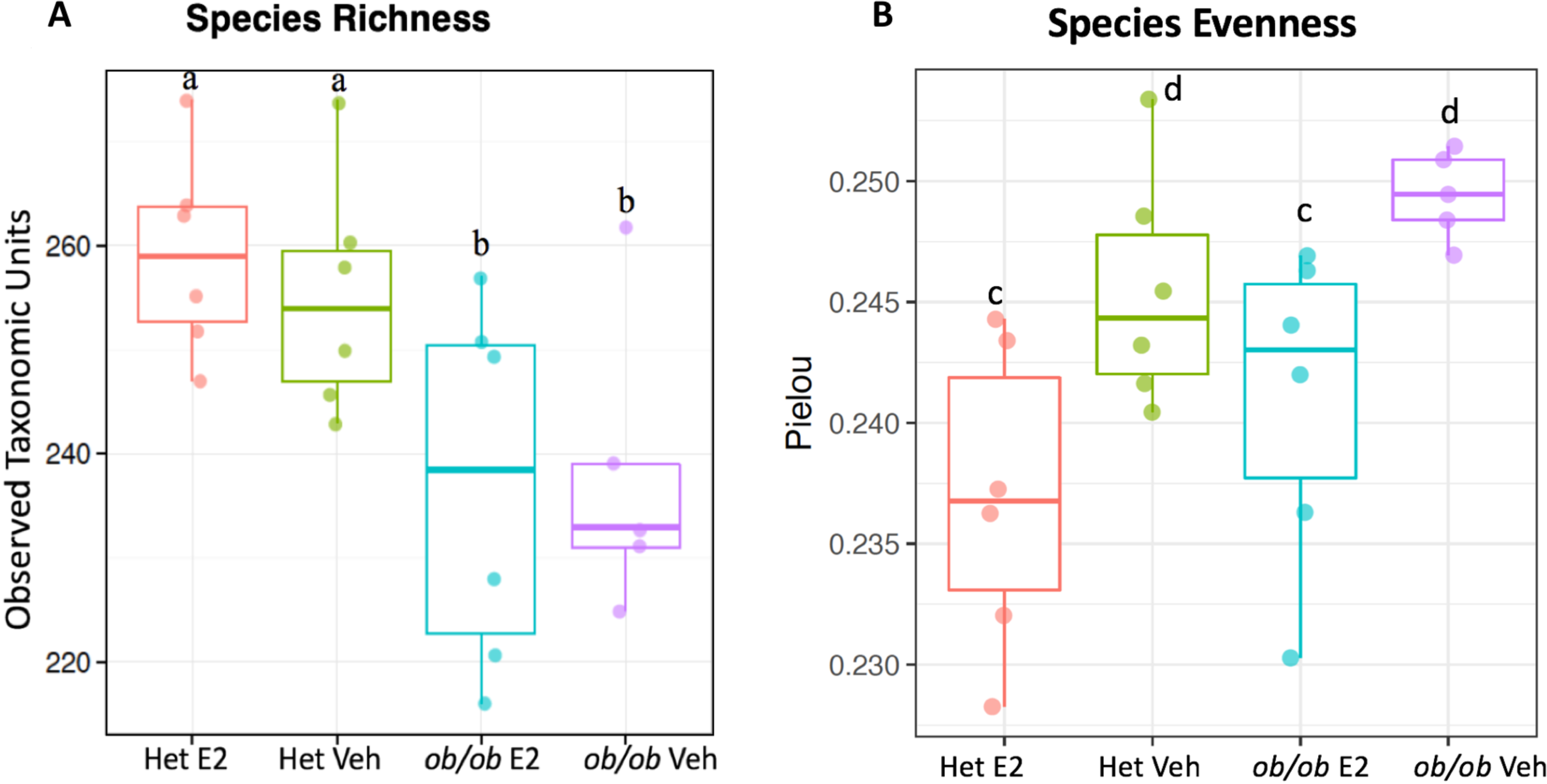
*ob/ob* mice have decreased species richness compared to controls, while estradiol treatment decreases species evenness. (A) *ob/ob* genotype is associated with lower species richness as measured by Observed Taxonomic Units. (B) E2 treatment is associated with lower species evenness as measured by the Pielou’s evenness index. “a” and “b” indicate groups with different species richness while “c” and “d” indicate groups with different species evenness (p<0.05, linear regression). Het E2 (n=6), Het Veh (n=6), *ob/ob* E2 (n=6) and *ob/ob* Veh (n=6).

The ordination plot based on Bray-Curtis Distance, which measures differences due to relative abundances and composition, showed distinct clustering of microbial communities from the four experimental groups (Figure 3; p=0.004, PERMANOVA). The first and second principal coordinates (PCo1 and PCo2) explain 29.4% and 17.5% of the variation in the microbial communities, respectively. Genotype, most likely through obesity, accounted for most of the difference observed as reflected by PCo1 (p=4E-8, t-test, Figure 3). There was also a clear effect of E2 treatment, represented by PCo2 (p=0.001, t-test, Figure 3).

**Figure 3.**
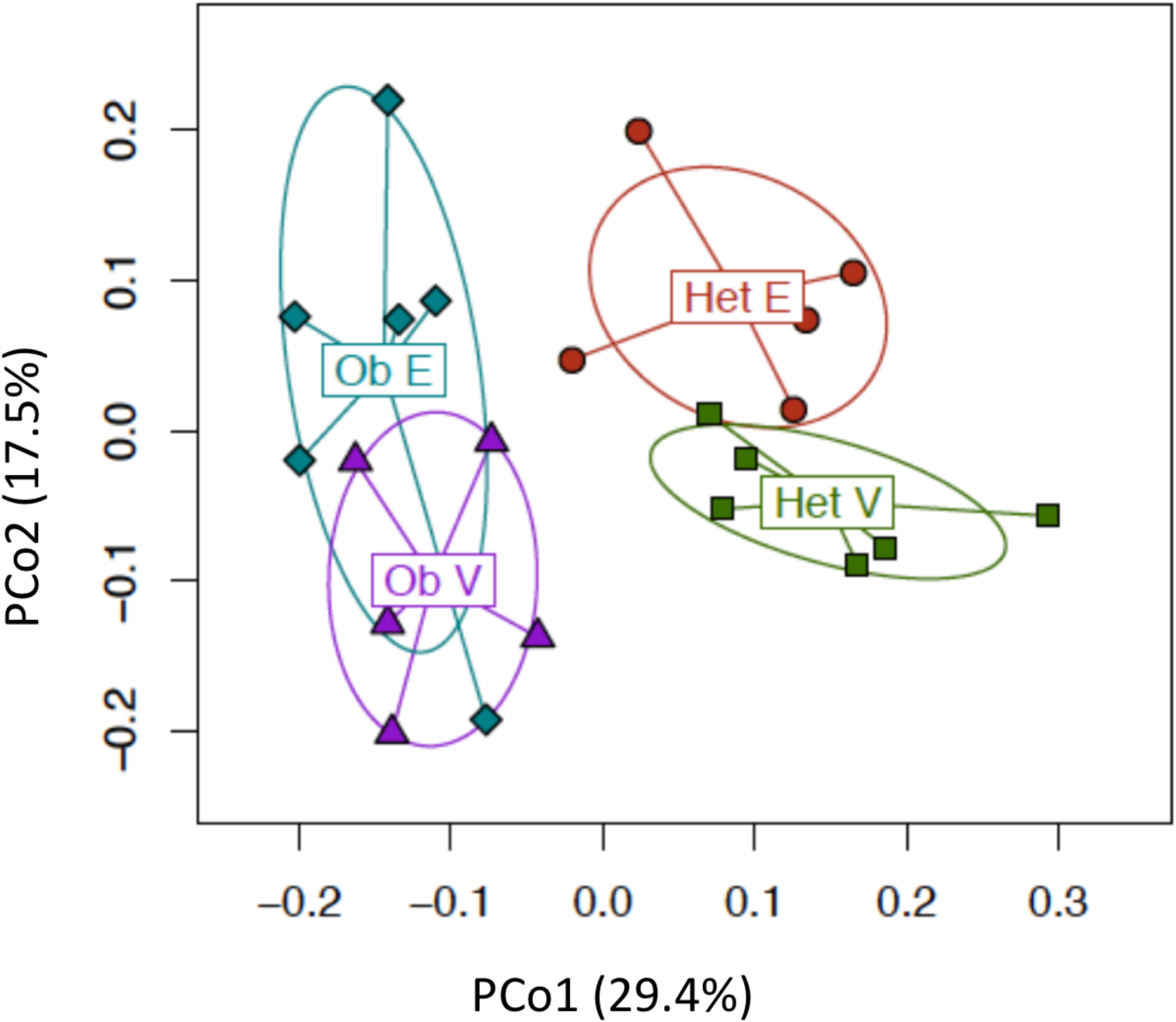
Gut microbial communities distinctly cluster as an effect of estradiol treatment and leptin. PCo1 and PCo2 clustering of each group. Bray-Curtis Distance was used to calculate principal coordinates 1 and 2 and PERMANOVA, to calculate the association with leptin or E2 on aggregate data over all days. Ob E = Estradiol-treated *ob/ob* (n=6), Ob V = Vehicle *ob/ob* (n=6), Het E= Estradiol-treated Het (n=6), and Het V= Vehicle Het mice (n=6).

Similar analyses were performed using Bray-Curtis Distance measures on each day to characterize the temporal changes in gut microbiota. The effect of E2 on β-diversity was robust and continuously observed from the 2^nd^ week of OVX (days 15-35). The effect of genotype was also present on days 7,15, 23 and 35, and a trend was observed on the remaining two days (4 and 31) (Table 1).

### Estradiol treatment alters relative abundances of intestinal microbiota

Gut microbial community composition was further analyzed on aggregated data from all days to identify the relative abundances at phylum, family and genus levels (Figure 4). E2-mediated shifts in relative abundances was evident on many of these taxa levels. In all four experimental groups, the most prevalent phyla (>90%) were Firmicutes, Bacteroidetes, Actinobacteria and Tenericutes (Figure 4A). The two most abundant families were S24-7 and Lachnospiraceae, within Bacteroidetes and Firmicutes, respectively (Figure 4B). At the genus level, *Coprococcus* and *Ruminococcus* (both within phylum Firmicutes) were the most abundant (Figure 4C).

To identify whether the differences in community structure were driven by differences in the relative abundances of particular microbial taxa, a differential abundance analysis based on overdispersed Poisson regression model was run on aggregate data to uncover the effects of treatment (Figure 5A) and genotype (Figure 5B). In the cladogram, the nodes in the first circle represent phyla, and the extending outer nodes in each level represent lower taxa within each phylum. A total of 26 taxa were differentially associated between E2 and Veh groups (Figure 6A). Firmicutes were more associated with Veh treatment, while Bacteroidetes were associated with E2 treatment (Figures 5A and 6A). Within Firmicutes, the class Erysipelotrichi, including *Allobaculum* and *Coprobacillus* spp., were more abundant in Veh than E2 mice. Similarly, the relative abundances of the class Bacilli, including its families Lactobacillaceae and Peptostreptococcaceae, were greater in Veh than E2 groups. Another family, Streptococcaceae, including *Lactococcus* spp., were also more associated with Veh mice. In contrast, the family Ruminococcaceae was more abundant in E2 mice. Within Bacteroidetes, the class Bacteroidia, and its lower taxa Bacteroidales and S24-7, were more abundant in E2 than Veh mice (Figures 5A and 6A).

**Figure 4.**
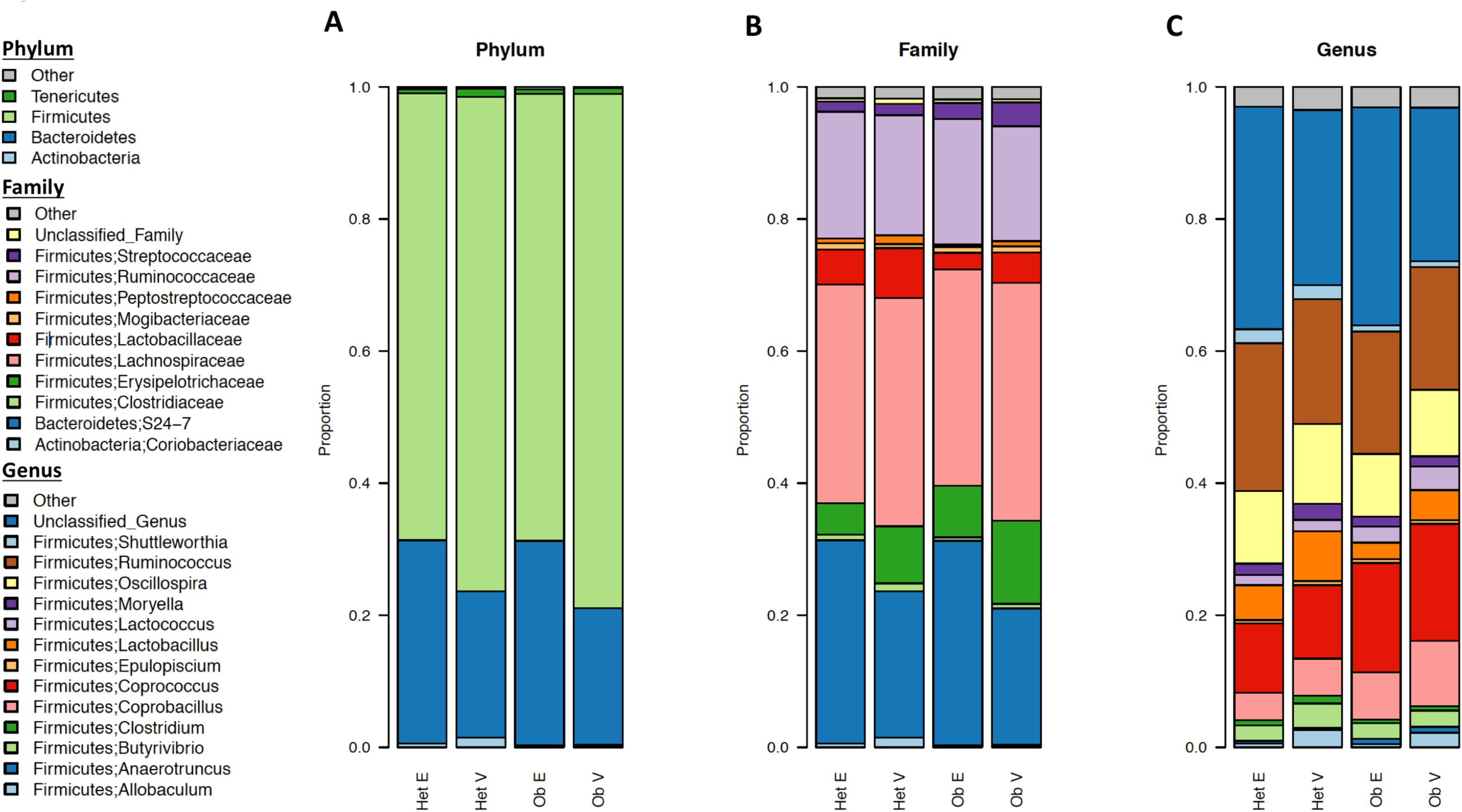
Gut microbiota associate with estradiol treatment and genotype at multiple taxa levels. Microbiota community structure at the (A) phylum, (B) family and (C) genus level, separated by treatment and genotype. Data are shown as relative proportion of the taxa identified. Taxa with prevalence of >10% or with a maximum proportion of > 0.2% were included. Ob E = Estradiol-treated *ob/ob* (n=6), Ob V = Vehicle *ob/ob* (n=6), Het E= Estradiol-treated Het (n=6), and Het V= Vehicle Het mice (n=6). An overdispersed Poisson regression was used to calculate associations of taxa with leptin and E2.

**Figure 5.**
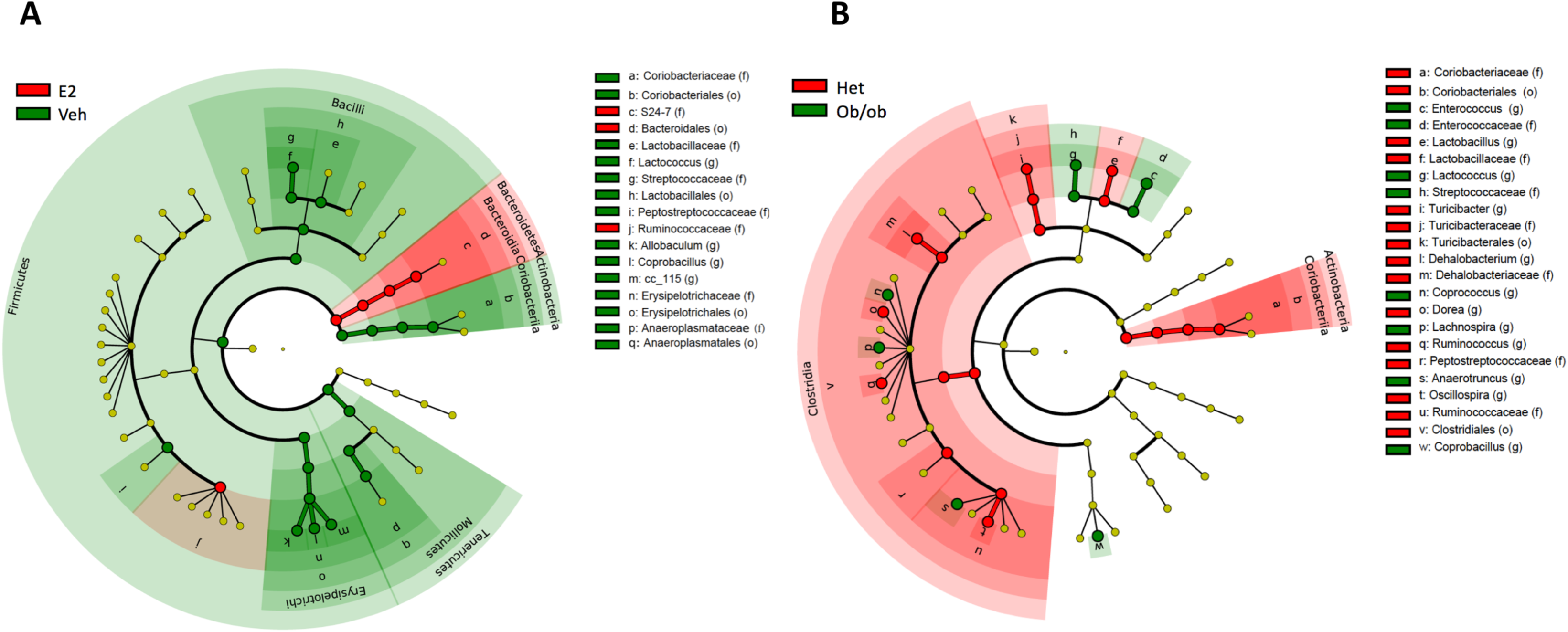
Estradiol and obesity alter the gut microbial composition. (A) Phyla and lower level taxa associated with estradiol (E2) or vehicle (Veh) treatment. (B) Phyla and lower level taxa associated with *ob/ob* or Het genotype. Het E2 (n=6), Het Veh (n=6), *ob/ob* E2 (n=6) and *ob/ob* Veh (n=6).

**Figure 6.**
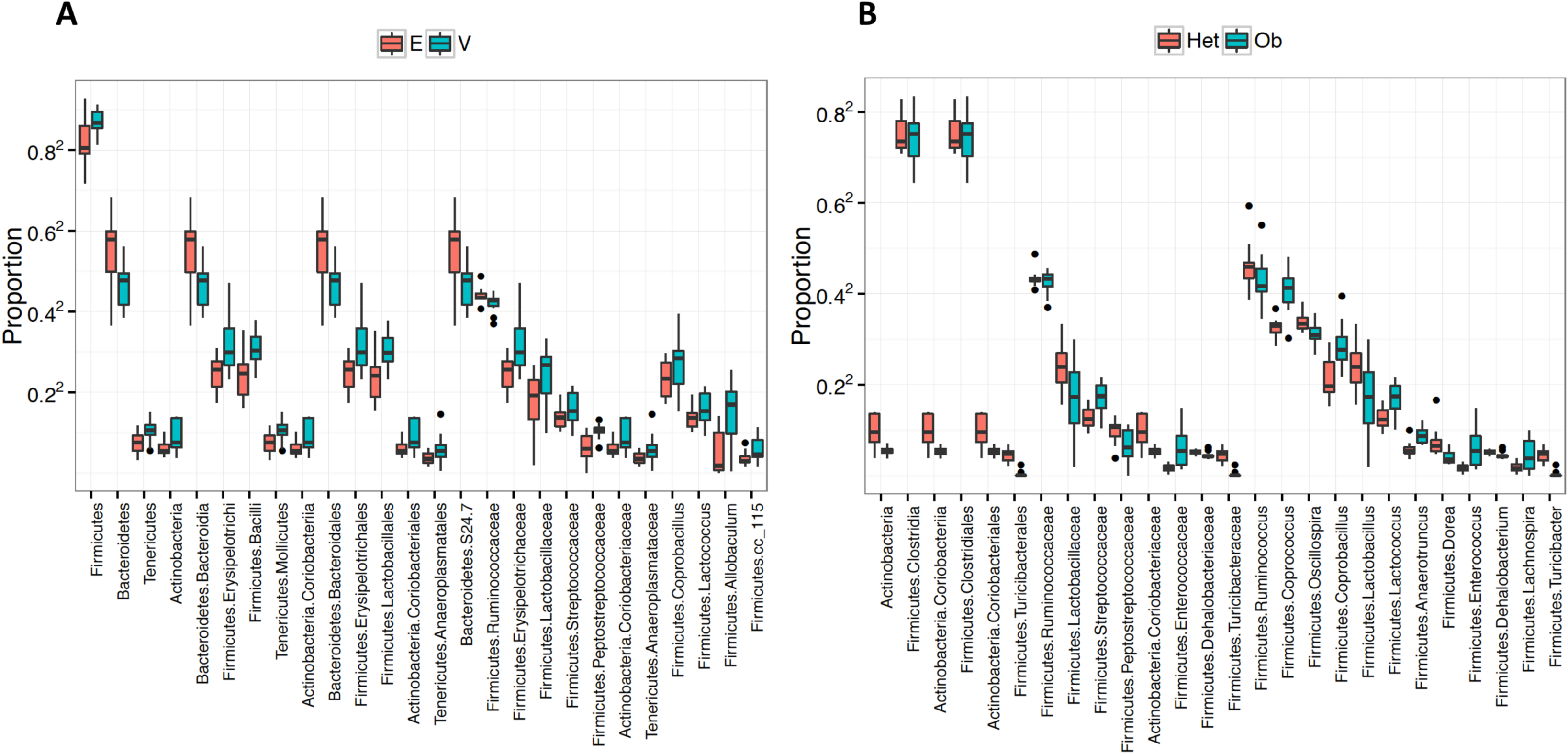
Gut microbiota associate with estradiol treatment and obesity at multiple taxa levels. Boxplots showing microbial taxa that are significantly altered by (A) E2 treatment and (B) genotype (overdispersed Poisson regression). Analysis was done on aggregate data from all 6 day points. Data are shown as relative proportion of the taxa identified. Taxa with prevalence of >10% or with a maximum proportion of > 0.2% were included. Estradiol-treated *ob/ob* (n=6), estradiol-treated Het (n=6), Ob V = vehicle *ob/ob* (n=6), Het V= vehicle Het mice (n=6).

For the taxa that differed across groups as detected from the aggregated data, the effect of E2 treatment on relative abundances was analyzed for each day. A comprehensive list of the taxa and associated q-values and fold changes between E2 and Veh mice for each day is provided in Supplemental Table 1. E2-treated mice resisted a decrease in relative abundances of Bacteroidetes compared to Veh mice on days 23 (q=0.007) and 31 (q=0.09) (Figure 7A). A main driver of the shifts in Bacteroidetes was the family S24-7. E2-treated mice resisted a decrease in S24-7 abundances compared to Veh mice on day 23 (q=0.015) (Figure 7B).

**Figure 7.**
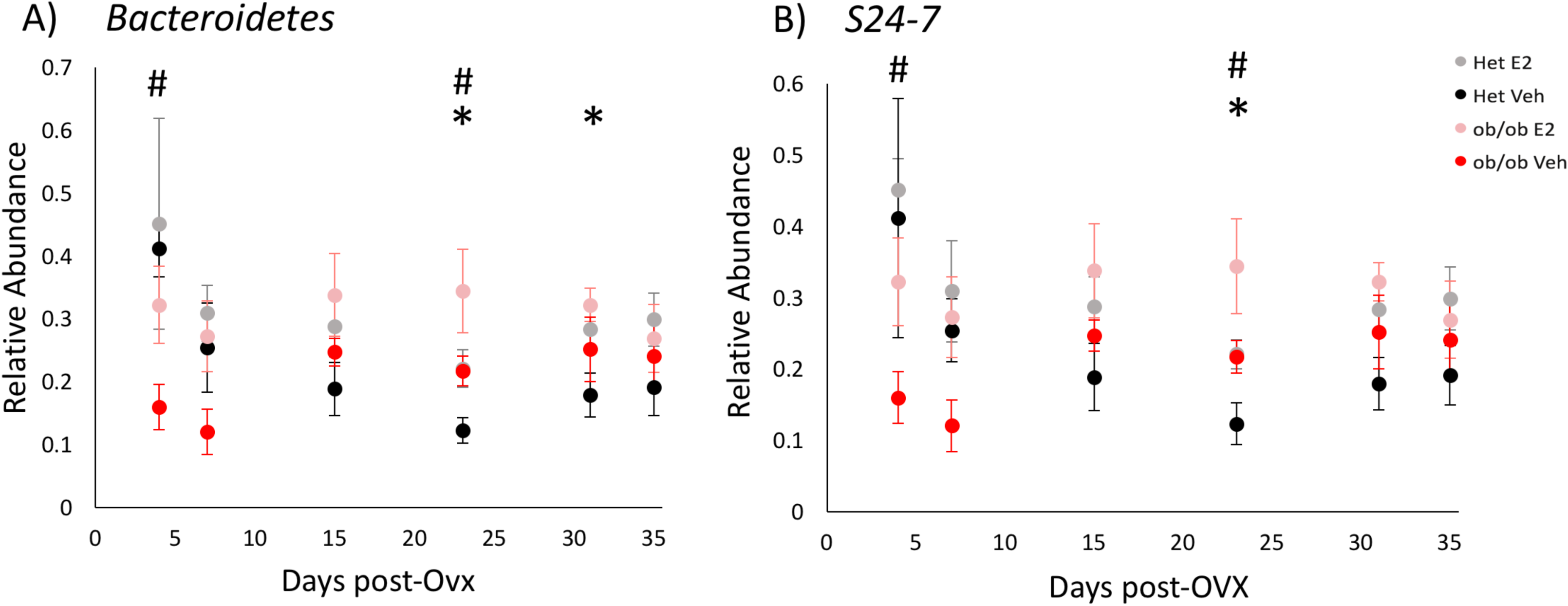
Estradiol and obesity alter the relative abundances of Bacteroidetes (phylum) and S24-7 (family) following the start of HFD. Relative abundances of the (A) the phylum *Bacteroidetes* and (B) its family *S24-7* over time. * indicates effects of E2 and # indicates effects of genotype (q<0.1, overdispersed Poisson regression). Het E2 (n=6), Het Veh (n=6), *ob/ob* E2 (n=6) and *ob/ob* Veh (n=6). Error bars indicate ±SEM.

Compared to Veh mice, E2-treated animals resisted changes in relative abundances of the phylum Firmicutes, with lower relative abundances of Firmicutes on days 23 (q=0.01), 31 (q=0.06) and 35 (q=0.04) compared to Veh mice (Figure 8A). Within Firmicutes, *Lactobacillus* spp. showed a significant reduction on day 31 (q=0.05) in E2-treated mice (Figure 8D).

**Figure 8.**
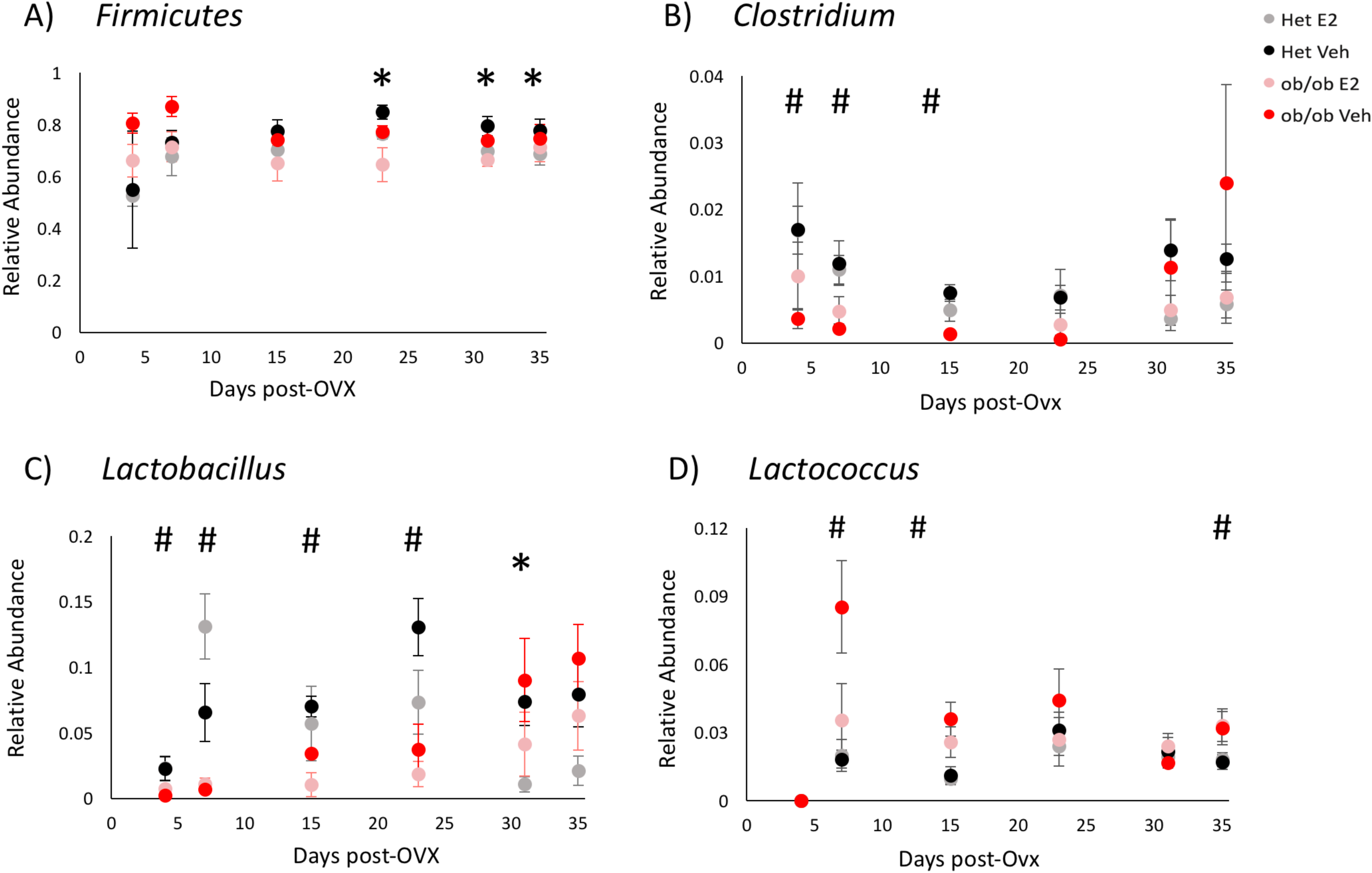
Estradiol and obesity alter the relative abundances of the phylum Firmicutes and its lower taxa. Relative abundances over time of the (A) phylum Firmicutes and genera (B) *Clostridium*, (C) *Lactobacillus* and (D) *Coprococcus*. * indicates effects of E2 and # indicates effects of genotype (q<0.1, overdispersed Poisson regression). Het E2 (n=6), Het Veh (n=6), *ob/ob* E2 (n=6) and *ob/ob* Veh (n=6). Error bars indicate ±SEM.

Estradiol also altered the relative abundance of the phylum Actinobacteria from days 15 to 35. On these days, E2 mice resisted an increase in the relative abundances of Actinobacteria compared to Veh mice (Figure 9A). The E2-mediated effect on Actinobacteria was due mostly to the family Coriobacteriaceae which resisted this increase also on days 15 (q=0.001), 23 (q<0.001) and 35 (q=0.03) (Figure 9B).

**Figure 9.**
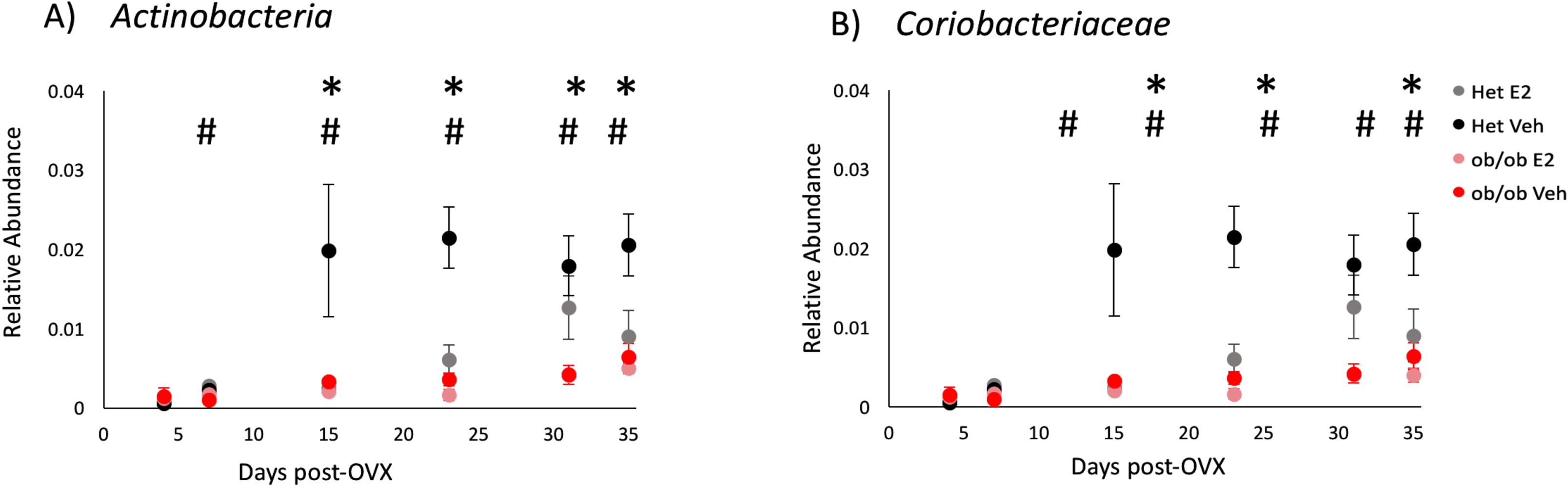
Estradiol and obesity resist increases in the relative abundances of the phylum Actinobacteria and its lower taxa. Relative abundances of the (A) phylum Actinobacteria and (B) family Coriobacteriaceae over time. * indicates effects of E2 and # indicates effects of genotype (q<0.1 overdispersed Poisson regression). Het E2 (n=6), Het Veh (n=6), *ob/ob* E2 (n=6) and *ob/ob* Veh (n=6). Error bars indicate ±SEM.

### Obesity modifies relative abundances of gut microbiota

As the gut microbial communities of the female mice clustered separately based on obesity (in *ob/ob* genotype) (Figure 3), we further analyzed the effects of obesity on relative abundances at the taxa levels (Figure 5B). A total of 26 identified taxa showed a differential association between *ob/ob* and Het mice (Figure 6B). Comparison of the gut microbial communities between genotypes revealed that Firmicutes, including its families Lactobacillaceae, Turibacteraceae, Peptostreptococcaceae, Dehalobacteriaceae, and Ruminococcaceae were more abundant in Veh Het mice than *ob/ob mice*. At the genus level, *Lactobacillus, Turibacter, Dehalobacterium, Dorea, Ruminococcus*, and *Oscillospira* were more abundant in control Het mice than *ob/ob* mice. Similarly, the phylum Actinobacteria and its family Coriobacteriaceae showed a greater abundance in control Het mice. In contrast, *Enterococcus, Coprococcus, Lachnospira, Anaerotruncus* and *Coprobacillus* spp. of the phylum Firmicutes were more abundant in *ob/ob* mice compared to Het mice (Figures 5B and 6B).

Differential abundance analysis across days showed that genotype affected the relative abundances of multiple taxa (Supplemental Table 2). Leptin increased the relative abundances of Bacteroidetes and its family S24-7 in Het mice at the beginning (day 4; q=0.04), but by day 23 (after 19 days on HFD; q=0.03), levels of these microbes were lower than in *ob/ob* mice (Figure 7). Within Firmicutes, *Clostridium, Lactobacillus* and *Lactococcus* spp. were altered by genotype. *Lactococcus* was more abundant in *ob/ob* mice on days 7 (q=0.01), 15 (q=0.002) and 35 (q=0.007) (Figure 8D). In contrast, *Clostridium* and *Lactobacillus* spp. were more abundant in Het mice. *Clostridium* was higher in Het mice on days 4 (q=0.04), 7 (q=0.01) and 15 (q<0.001) than *ob/ob* mice (Figure 8B). Similarly, the relative abundance of *Lactobacillus* increased in Het mice compared to *ob/ob* mice on days 4 (q=0.01), 7 (q<0.001) and 23 (q=0.001) (Figure 8D).

Obesity also altered the relative abundance of the phylum Actinobacteria over time. Compared to *ob/ob* mice, Het mice had an increase in the relative abundance of Actinobacteria from days 7 to 35 (Figure 9A). This leptin-dependent increase in Actinobacteria in Hets was due mostly to Coriobacteriaceae (family), also from days 7 to 35 (Figure 9B), suggesting this microbial family is influenced by leptin or obesity status during HFD.

A subset of microbial taxa was affected by both E2 and leptin. In particular, Streptococcaceae, *Lactococcus* and *Coprobacillus* were decreased, whereas Ruminococcaceae were increased by E2 and leptin. In addition, these two hormones exerted opposing effects on other families, such that E2 decreased, and leptin (in Het controls) increased, abundances of Coriobacteriaceae, Lactobacillaceae and Peptostreptococcaceae.

## Discussion

Using leptin-deficient *ob/ob* female mice fed a HFD, we investigated the effects of estradiol and obesity (due to leptin deficiency) on body weight, energy intake, and gut microbiota in the present study. We found that E2 treatment decreases food intake and protects against HFD-induced weight gain in both *ob/ob* (obese) and Het (lean) females. *ob/ob* mice had increased HFD intake and greater body weight, compared to the Het controls. The differences in food intake and body weight between the two genotypes are reflected in their changes in gut microbiota. Both E2 treatment and genotype altered gut microbiota diversity. o*b/ob* female mice exhibited lower species richness, which supports previous studies that link obesity with a reduced gut microbiota diversity in humans and male mice [60-62]. Interestingly, E2 treatment was associated with lower species evenness compared to Veh mice. At the phylum level, E2 treatment slowed down a HFD-induced decrease in Bacteroidetes and increases in Firmicutes and Actinobacteria compared to Veh controls. Leptin also altered the relative abundances of Bacteroidetes and Firmicutes and profoundly increased Actinobacteria. Many taxa were also affected by both leptin and E2. These findings suggest that E2 and leptin can act independently, or interact together, to modulate gut microbiota to mediate energy regulation during HFD intake in females.

### E2 alters gut microbial community diversity in ob/ob female mice on a HFD

We and others have shown that E2 protects female mice from HFD-induced obesity [10, 11, 13, 63-66]. Estrogens exert protective effects by acting directly on brain [10-12, 67], pancreas [68], liver [63, 64, 66, 69], adipose tissue [66], and muscle [70, 71] to regulate energy production and utilization. Moreover, long-term E2 treatment improves glucose tolerance and insulin sensitivity and attenuates lipid synthesis in liver in *ob/ob* female mice [72]. The present findings provide evidence that the modulation of gut microbiota is another mechanism by which E2 mediates energy homeostasis. While E2 was associated with a decrease in the gut microbial evenness in the current study, an increase in microbial diversity has been reported in cycling rats and E2-treated female mice [38, 73]. These differences across studies could be due to fluctuating estrogens and other ovarian hormones in the cycling rats and/or species differences [38], and a much higher dose of estradiol (2.5 mg/day, consistent with levels at pregnancy) used in the mice [73]. Alternatively, while a lower microbial diversity is usually associated with obesity and metabolic disorders [74, 75], a change in relative abundance without any changes in microbial richness or evenness can also alter microbial homeostasis [45].

### Potential implications of E2-mediated alterations in gut microbiota composition in ob/ob female mice on HFD

E2 attenuated a longitudinal shift in the two major phyla Bacteroidetes and Firmicutes, compared to the Veh controls. In particular, the order Bacteroidales, and its families S24-7 and Ruminococcaceae were positively associated with E2 treatment, whereas *Allobaculum* spp. were negatively associated. The current and previous studies suggest that various gut microbial communities collectively exert E2-mediated protection against HFD-induced weight gain [38, 43].

Functional predictions of microbial taxa in a population are mostly derived from the gene families they express. S24-7 and Ruminococcaceae, both microbial families upregulated by E2 in the current study, produce short chain fatty acids (SCFA; fermented products of dietary fibers) that protect against inflammation [76-78]. In particular, SCFA, including butyrate, maintain a low pH in the gastrointestinal tract and aid in nutrient absorption and pathogen inactivation [79]. Similarly, both S24-7 and Ruminococcaceae are downregulated in inflammatory conditions, including Crohn’s disease, colitis and type I diabetes [80-82]. Taken together, the current findings suggest that E2 provides protection against diet-induced metabolic disorders in females by maintaining healthy levels of S24-7 and Ruminococcaceae.

Firmicutes, altered by E2 in the current study, is generally associated with obesity in rodents and humans [83]. Studies in males have shown that a high caloric diet profoundly affects Firmicutes [29, 84, 85]. Consistent with these findings in males, in the present study, Veh female mice on a HFD gained weight and had an increase in Firmicutes. However, E2-treated mice resisted this HFD-induced increase, suggesting that modulation of these microbes contributes to the regulation of energy homeostasis by estrogens. Firmicutes contain more OTUs that are efficient energy producers than Bacteroidetes, leading to increased calorie absorption and weight gain [86]. Additionally, some members of Firmicutes promote lipid droplets formation increasing fatty acid absorption and weight gain [87]. In the current study, E2 altered relative abundances of Firmicutes in both Het and *ob/ob* mice, suggesting that E2 affects this phylum independent of body weight and leptin levels.

### Obesity alters gut microbial community diversity in female mice on a HFD

The obese leptin-deficient (*ob/ob*) mice had less diverse microbiota as observed by species richness, suggesting that leptin is required for an enriched microbial ecosystem. In support, obesity is associated with a decreased microbial diversity in humans [60]. Furthermore, a Danish study on men and women found that higher levels of leptin in obese populations were associated with a lower richness of the gut microbial communities, suggesting that optimum levels of leptin are associated with metabolic health and a more diverse gut microbiota [61]. Taken together, the present results indicate that a shift in leptin levels in either direction disrupts metabolic and microbial homeostasis.

### Potential implications of obesity-induced alterations in gut microbiota composition in female mice on a HFD

While we observed a positive correlation of Bacteroidetes and its family S24-7 with leptin (in Het mice) at the beginning of the study, by day 23, their levels dropped to even lower levels than *ob/ob* mice, suggesting a pronounced negative effect of HFD on Bacteroidetes in Hets. Within Firmicutes, *ob/ob* mice had lower relative abundances of *Clostridium* and *Lactobacillus* compared to control Het mice. In support, high leptin levels associate with Lactobacillales, the order containing *Lactobacillus* spp., in African-American men [88]. Most *Lactobacilli* spp. are negatively correlated with inflammation and exert their beneficial effects through modulation of the immune system via macrophage and dendritic cell functions [89, 90]. These findings suggest that regulation of gut microbiota by leptin is an important mechanism for metabolic homeostasis.

While *ob/ob* mice weighed almost twice that of Het mice in the beginning, the percent weight gain was greater in Veh Hets compared to Veh *ob/ob* mice. Interestingly, the phylum Actinobacteria showed a greater relative abundance in Hets compared to ob/ob mice, primarily due to an increase in its family Coriobacteraceae. Actinobacteria is increased in obese humans and is associated with ulcerative colitis [60, 75]. More than two-thirds of the obesity-related human gut microbes belong to Actinobacteria [60, 91]. In particular, Coriobacteriaceae have been positively associated with increased cholesterol absorption in hamsters and obese humans [92, 93]. A profound increase in the abundance of Coriobacteriaceae in Veh Het mice, the group that showed the highest percent weight gain, but not in E2 Hets, suggests this OTU is associated with HFD-induced weight gain.

### Possible mechanisms for estradiol- and leptin-regulation of gut microbiota

Estrogens elicit many effects on physiology and behavior by binding to their intracellular and membrane receptors [94-97]. While estrogen receptor-α (ERα) and ERβ have been implicated in the effects of estrogens on metabolism [98-100], ERα appears to be the primary contributor to energy balance [98, 101-103]. ERα knock-out mice exhibit increased visceral adiposity, impaired glucose tolerance and elevated insulin levels [103]. Furthermore, systemic activation of ERα, but not ERβ, decreases food intake and body weight in female rats [102]. While the effects of ERα on gut microbiota have not been investigated directly, a study using ERβ knock-out female mice suggests that ERβ influences gut microbiota in a diet-specific manner [104]. In support of this finding, ERβ is expressed in colon epithelial cells [105, 106]. Alternatively, given that estrogens reduce food intake [10], it is also possible that E2 influences gut microbiota by altering nutrition availability.

Estrogens also act as direct substrates for gut microbiota. For example, microbes with β-glucuronidase and β-glucosidase enzymes, including *Lactobacillus, Bifidobacterium*, and *Clostridium* spp., convert inactive estrogens into their active forms through deconjugation [107-109]. Estrogens can also alter metabolic outcome and immune response through direct actions on gut microbiota. For example, production of lipopolysaccharides (LPS), an endotoxin produced by Gram-negative bacteria, is elevated after 4 weeks of HFD and is associated with obesity and insulin resistance in male mice [110, 111]. E2 treatment may protect from HFD-induced metabolic disorders through blocking LPS activation [112]. Additionally, E2 upregulates intestinal alkaline phosphatase, a protein with protective role in the gut epithelium through attenuation of proinflammatory signals in the intestine [43]. Taken together with the present findings, estrogens can alter gut microbiota through multiple direct and indirect mechanisms to protect from metabolic disorders.

The mechanisms by which leptin affects gut microbiota are not well understood. Intestinal epithelium expresses leptin receptors (LepR), but ablation of these receptors does not alter gut microbiota or body weight in male mice [113]. In females, however, leptin could act in concert with ERß in the gut epithelium [105, 106]. While genotype effects may be primarily dependent on leptin, it is important to note that *ob/ob* mice are obese and Het mice are lean. Obesity is an independent modulator of gut microbiota. Obese and lean twins exhibit differences in gut microbial diversity [60], and the obese phenotype can be transferred to lean recipients through the transplant of gut microbiota from obese donors [29, 30]. In addition, female *ob/ob* mice have reduced estrogen levels and are sterile, and thus have altered hormone-dependent development [26, 114]. These findings suggest that leptin affects host physiology, including gut microbial homeostasis through multiple mechanisms.

This study provides critical insights into the roles of E2 and leptin in HFD-induced obesity and gut microbiota. While the identification of taxa that differ between treatments is the first step towards defining their functions, many microbes share redundant metabolic functions [74]. Metagenomic analysis, or correlation analysis between gut microbiota and metabolic changes, will provide further important insights on microbial community-specific functions and their effects on metabolism.

## Conclusion

This study provides evidence for the function of estrogens and leptin in energy homeostasis through novel actions on gut microbiota. Estrogens altered the microbiota community in female mice that were fed a HFD. Furthermore, the use of female *ob/ob* mice that lack leptin suggests that leptin regulates gut microbiota in females. Given that many metabolic and inflammatory bowel diseases are characterized by disturbances in the gut microbiome, understanding the functions of E2 and leptin in modulating gut microbiome in females has important therapeutic applications for women with metabolic disorders.

## Acknowledgements

This work was funded by NIH R01 DK61935 (MJT) and a Re-Entry award (EPB) from the National Institute of Diabetes and Digestive and Kidney Diseases. The authors thank Dr. Cassandra Pattanayak, Director of the Quantitative Analysis Institute at Wellesley College, for input on statistical analysis.

## References

1. Asarian L, Geary N: Modulation of appetite by gonadal steroid hormones. Philos Trans R Soc Lond B Biol Sci 2006, 361(1471):1251–1263.

2. Wade GN, Gray JM: Gonadal effects on food intake and adiposity: a metabolic hypothesis. Physiol Behav 1979, 22(3):583–593.

3. Clegg DJ: Minireview: the year in review of estrogen regulation of metabolism. Mol Endocrinol 2012, 26(12):1957–1960.

4. McCarthy MM: Estradiol and the developing brain. Physiol Rev 2008, 88(1):91–124.

5. Pfaff D, Waters E, Khan Q, Zhang X, Numan M: Minireview: estrogen receptor-initiated mechanisms causal to mammalian reproductive behaviors. Endocrinology 2011, 152(4):1209–1217.

6. Blaustein JD: Neuroendocrine regulation of feminine sexual behavior: lessons from rodent models and thoughts about humans. Annu Rev Psychol 2008, 59:93–118.

7. Roquer J, Campello AR, Gomis M: Sex differences in first-ever acute stroke. Stroke 2003, 34(7):1581–1585.

8. Carr MC: The emergence of the metabolic syndrome with menopause. J Clin Endocrinol Metab 2003, 88(6):2404–2411.

9. Guo SS, Zeller C, Chumlea WC, Siervogel RM: Aging, body composition, and lifestyle: the Fels Longitudinal Study. Am J Clin Nutr 1999, 70(3):405–411.

10. Bless EP, Reddy T, Acharya KD, Beltz BS, Tetel MJ: Oestradiol and diet modulate energy homeostasis and hypothalamic neurogenesis in the adult female mouse. J Neuroendocrinol 2014, 26(11):805–816.

11. Bless EP, Yang J, Acharya KD, Nettles SA, Vassoler FM, Byrnes EM, Tetel MJ: Adult neurogenesis in the female mouse hypothalamus: Estradiol and high-fat diet alter the generation of newborn neurons expressing estrogen receptor alpha. eNeuro 2016, 3(4):1–11.

12. Brown LM, Clegg DJ: Central effects of estradiol in the regulation of food intake, body weight, and adiposity. J Steroid Biochem Mol Biol 2010, 122(1-3):65–73.

13. Xu Y, Nedungadi TP, Zhu L, Sobhani N, Irani BG, Davis KE, Zhang X, Zou F, Gent LM, Hahner LD et al: Distinct hypothalamic neurons mediate estrogenic effects on energy homeostasis and reproduction. Cell Metab 2011, 14(4):453–465.

14. Erickson JC, Hollopeter G, Palmiter RD: Attenuation of the obesity syndrome of ob/ob mice by the loss of neuropeptide Y. Science 1996, 274(5293):1704–1707.

15. Vaisse C, Halaas JL, Horvath CM, Darnell JE, Jr., Stoffel M, Friedman JM: Leptin activation of Stat3 in the hypothalamus of wild-type and ob/ob mice but not db/db mice. Nat Genet 1996, 14(1):95–97.

16. Sahu A: Leptin signaling in the hypothalamus: emphasis on energy homeostasis and leptin resistance. Frontiers in neuroendocrinology 2003, 24(4):225–253.

17. Green ED, Maffei M, Braden VV, Proenca R, DeSilva U, Zhang Y, Chua SC, Jr., Leibel RL, Weissenbach J, Friedman JM: The human obese (OB) gene: RNA expression pattern and mapping on the physical, cytogenetic, and genetic maps of chromosome 7. Genome research 1995, 5(1):5–12.

18. Coleman DL: Diabetes-Obesity Syndromes in Mice. Diabetes 1982, 31:1–6.

19. Pelleymounter MA, Cullen M.J., Baker, M.B.,, Hecht R, Winters, D., Boone, T., Collins, F.: Effects of the obese Gene Product on Body Weight Regulation in ob/ob Mice. Science 1995, 269:540–543.

20. Mistry AM, Andrew G. Swick AG, Romsos DR: Leptin Rapidly Lowers Food Intake and Elevates Metabolic Rates in Lean and ob/ob Mice. J Nutr 1997, 127(10):2065–2072.

21. Shimomura K, Shimizu H, Tsuchiya T, Abe Y, Uehara Y, Mori M: Is leptin a key factor which develops obesity by ovariectomy? Endocr J 2002, 49(4):417–423.

22. Pelleymounter MA, Cullen MJ, Baker MB, Hecht R, Winters D, Boone T, Collins F: Effects of the obese gene product on body weight regulation in ob/ob mice. Science 1995, 269(5223):540–543.

23. Steppan CM, Crawford DT, Chidsey-Frink KL, Ke H, Swick AG: Leptin is a potent stimulator of bone growth in ob/ob mice. Regul Pept 2000, 92(1-3):73–78.

24. Clegg DJ, Riedy CA, Smith KA, Benoit SC, Woods SC: Differential sensitivity to central leptin and insulin in male and female rats. Diabetes 2003, 52(3):682–687.

25. Clegg DJ, Brown LM, Woods SC, Benoit SC: Gonadal hormones determine sensitivity to central leptin and insulin. Diabetes 2006, 55(4):978–987.

26. Chehab FF, Lim ME, Lu R: Correction of the sterility defect in homozygous obese female mice by treatment with the human recombinant leptin. Nat Genet 1996, 12(3):318–320.

27. Soto M, Herzog C, Pacheco JA, Fujisaka S, Bullock K, Clish CB, Kahn CR: Gut microbiota modulate neurobehavior through changes in brain insulin sensitivity and metabolism. Mol Psychiatry 2018.

28. Cani PD: Human gut microbiome: hopes, threats and promises. Gut 2018, 67(9):1716–1725.

29. Turnbaugh PJ, Ridaura VK, Faith JJ, Rey FE, Knight R, Gordon JI: The effect of diet on the human gut microbiome: a metagenomic analysis in humanized gnotobiotic mice. Sci Transl Med 2009, 1:6ra14.

30. Ridaura VK, Faith JJ, Rey FE, Cheng J, Duncan AE, Kau AL, Griffin NW, Lombard V, Henrissat B, Bain JR et al: Gut microbiota from twins discordant for obesity modulate metabolism in mice. Science 2013, 341(6150):1241214.

31. Ellekilde M, Selfjord E, Larsen CS, Jakesevic M, Rune I, Tranberg B, Vogensen FK, Nielsen DS, Bahl MI, Licht TR et al: Transfer of gut microbiota from lean and obese mice to antibiotic-treated mice. Sci Rep 2014, 4:5922.

32. Ley RE, Hamady M, Lozupone C, Turnbaugh P, Ramey RR, Bircher JS, Schlegel ML, Tucker TA, Schrenzel MD, Knight R et al: Evolution of mammals and their gut microbes. Science 2008, 320(5883):1647–1651.

33. Round JL, Mazmanian SK: The gut microbiome shapes intestinal immune responses during health and disease. Nature reviews Immunology 2009, 9(5):313–323.

34. Kang C, Wang B, Kaliannan K, Wang X, Lang H, Hui S, Huang L, Zhang Y, Zhou M, Chen M et al: Gut Microbiota Mediates the Protective Effects of Dietary Capsaicin against Chronic Low-Grade Inflammation and Associated Obesity Induced by High-Fat Diet. mBio 2017, 8(3).

35. Clarke G, Stilling RM, Kennedy PJ, Stanton C, Cryan JF, Dinan TG: Minireview: Gut microbiota: the neglected endocrine organ. Mol Endocrinol 2014, 28(8):1221–1238.

36. Markle JG, Frank DN, Mortin-Toth S, Robertson CE, Feazel LM, Rolle-Kampczyk U, von Bergen M, McCoy KD, Macpherson AJ, Danska JS: Sex differences in the gut microbiome drive hormone-dependent regulation of autoimmunity. Science 2013, 339(6123):1084–1088.

37. Tetel MJ, de Vries GJ, Melcangi RC, Panzica G, O’Mahony SM: Steroids, Stress, and the Gut Microbiome-Brain Axis. J Neuroendocrinol 2017.

38. Moreno-Indias I, Sanchez-Alcoholado L, Sanchez-Garrido MA, Martin-Nunez GM, Perez-Jimenez F, Tena-Sempere M, Tinahones FJ, Queipo-Ortuno MI: Neonatal Androgen Exposure Causes Persistent Gut Microbiota Dysbiosis Related to Metabolic Disease in Adult Female Rats. Endocrinology 2016, 157(12):4888–4898.

39. Org E, Mehrabian M, Parks BW, Shipkova P, Liu X, Drake TA, Lusis AJ: Sex differences and hormonal effects on gut microbiota composition in mice. Gut microbes 2016:0.

40. Foster JA, McVey Neufeld KA: Gut-brain axis: how the microbiome influences anxiety and depression. Trends Neurosci 2013, 36(5):305–312.

41. Jasarevic E, Morrison KE, Bale TL: Sex differences in the gut microbiome-brain axis across the lifespan. Philos Trans R Soc Lond B Biol Sci 2016, 371(1688):20150122.

42. Mueller S, Saunier K, Hanisch C, Norin E, Alm L, Midtvedt T, Cresci A, Silvi S, Orpianesi C, Verdenelli MC et al: Differences in fecal microbiota in different European study populations in relation to age, gender, and country: a cross-sectional study. Appl Environ Microbiol 2006, 72(2):1027–1033.

43. Kaliannan K, Robertson RC, Murphy K, Stanton C, Kang C, Wang B, Hao L, Bhan AK, Kang JX: Estrogen-mediated gut microbiome alterations influence sexual dimorphism in metabolic syndrome in mice. Microbiome 2018, 6(1):205.

44. Yurkovetskiy L, Burrows M, Khan AA, Graham L, Volchkov P, Becker L, Antonopoulos D, Umesaki Y, Chervonsky AV: Gender bias in autoimmunity is influenced by microbiota. Immunity 2013, 39(2):400–412.

45. Ley RE, Backhed F, Turnbaugh P, Lozupone CA, Knight RD, Gordon JI: Obesity alters gut microbial ecology. Proc Natl Acad Sci U S A 2005, 102(31):11070–11075.

46. Murphy EF, Cotter PD, Healy S, Marques TM, O’Sullivan O, Fouhy F, Clarke SF, O’Toole PW, Quigley EM, Stanton C et al: Composition and energy harvesting capacity of the gut microbiota: relationship to diet, obesity and time in mouse models. Gut 2010, 59(12):1635–1642.

47. Ley RE, Turnbaugh PJ, Klein S, Gordon JI: Microbial ecology: human gut microbes associated with obesity. Nature 2006, 444(7122):1022–1023.

48. Yang M, Liu Y, Xie H, Wen Z, Zhang Y, Wu C, Huang L, Wu J, Xie C, Wang T et al: Gut Microbiota Composition and Structure of the Ob/Ob and Db/Db Mice. Int J Endocrinol 2019, 2019:1394097.

49. Chung WK, Belfi K, Chua M, Wiley J, Mackintosh RD, Nicolson M, Boozer CN, Leibel RL: Heterozygosity for Lepob or Leprdb affects body composition and leptin homeostasis in adult mice. Am J Physiol 1998.

50. Caporaso JG, Lauber CL, Walters WA, Berg-Lyons D, Lozupone CA, Turnbaugh PJ, Fierer N, Knight R: Global patterns of 16S rRNA diversity at a depth of millions of sequences per sample. Proc Natl Acad Sci U S A 2011, 108 Suppl 1:4516–4522.

51. Chen X, Johnson S, Jeraldo P, Wang J, Chia N, Kocher JA, Chen J: Hybrid-denovo: a de novo OTU-picking pipeline integrating single-end and paired-end 16S sequence tags. GigaScience 2018, 7(3):1–7.

52. Cole JR, Wang Q, Fish JA, Chai B, McGarrell DM, Sun Y, Brown CT, Porras-Alfaro A, Kuske CR, Tiedje JM: Ribosomal Database Project: data and tools for high throughput rRNA analysis. Nucleic Acids Res 2014, 42(Database issue):D633–642.

53. Price MN, Dehal PS, Arkin AP: FastTree: computing large minimum evolution trees with profiles instead of a distance matrix. Mol Biol Evol 2009, 26(7):1641–1650.

54. Pielou EC: The measurement of diversity in different types of biological collections. J Theor Biol 1966, 13:131–144.

55. McArdle BH, Anderson MJ: Fitting multivariate models to community data: a comment on distance-based redundancy analysis. Ecology 2001, 82(1):290–297.

56. Chen J, Toyomasu Y, Hayashi Y, Linden DR, Szurszewski JH, Nelson H, Farrugia G, Kashyap PC, Chia N, Ordog T: Altered gut microbiota in female mice with persistent low body weights following removal of post-weaning chronic dietary restriction. Genome Med 2016, 8(1):103.

57. Chen L, Reeve J, Zhang L, Huang S, Wang X, Chen J: GMPR: A robust normalization method for zero-inflated count data with application to microbiome sequencing data. PeerJ 2018, 6:e4600.

58. Benjamini Y, Hochberg Y: Controlling the False Discovery Rate: A Practical and Powerful Approach to MultipleTesting. J R Statist Soc B (Methodological) 1995, 57:289–300.

59. Asnicar F, Weingart G, Tickle TL, Huttenhower C, Segata N: Compact graphical representation of phylogenetic data and metadata with GraPhlAn. PeerJ 2015, 3:e1029.

60. Turnbaugh PJ, Hamady M, Yatsunenko T, Cantarel BL, Duncan A, Ley RE, Sogin ML, Jones WJ, Roe BA, Affourtit JP et al: A core gut microbiome in obese and lean twins. Nature 2009, 457(7228):480–484.

61. Le Chatelier E, Nielsen T, Qin J, Prifti E, Hildebrand F, Falony G, Almeida M, Arumugam M, Batto JM, Kennedy S et al: Richness of human gut microbiome correlates with metabolic markers. Nature 2013, 500(7464):541–546.

62. Ravussin Y, Koren O, Spor A, LeDuc C, Gutman R, Stombaugh J, Knight R, Ley RE, Leibel RL: Responses of gut microbiota to diet composition and weight loss in lean and obese mice. Obesity (Silver Spring, Md) 2012, 20(4):738–747.

63. Camporez JP, Jornayvaz FR, Lee HY, Kanda S, Guigni BA, Kahn M, Samuel VT, Carvalho CR, Petersen KF, Jurczak MJ et al: Cellular mechanism by which estradiol protects female ovariectomized mice from high-fat diet-induced hepatic and muscle insulin resistance. Endocrinology 2013, 154(3):1021–1028.

64. Riant E, Waget A, Cogo H, Arnal JF, Burcelin R, Gourdy P: Estrogens protect against high-fat diet-induced insulin resistance and glucose intolerance in mice. Endocrinology 2009, 150(5):2109–2117.

65. Mamounis KJ, Hernandez MR, Margolies N, Yasrebi A, Roepke TA: Interaction of 17beta-estradiol and dietary fatty acids on energy and glucose homeostasis in female mice. Nutr Neurosci 2017:1–14.

66. Bryzgalova G, Lundholm L, Portwood N, Gustafsson JA, Khan A, Efendic S, Dahlman-Wright K: Mechanisms of antidiabetogenic and body weight-lowering effects of estrogen in high-fat diet-fed mice. Am J Physiol Endocrinol Metab 2008, 295(4):E904–912.

67. Gao Q, Horvath TL: Neurobiology of feeding and energy expenditure. Annu Rev Neurosci 2007, 30:367–398.

68. Tiano JP, Mauvais-Jarvis F: Molecular mechanisms of estrogen receptors’ suppression of lipogenesis in pancreatic beta-cells. Endocrinology 2012, 153(7):2997–3005.

69. Yasrebi A, Rivera JA, Krumm EA, Yang JA, Roepke TA: Activation of Estrogen Response Element-Independent ERalpha Signaling Protects Female Mice From Diet-Induced Obesity. Endocrinology 2017, 158(2):319–334.

70. Ribas V, Drew BG, Zhou Z, Phun J, Kalajian NY, Soleymani T, Daraei P, Widjaja K, Wanagat J, et al.: Skeletal muscle action of estrogen receptor a is critical for the maintenance of mitochondrial function andmetabolic homeostasis in females. Sci Transl Med 2016.

71. Campbell SE, Mehan KA, Tunstall RJ, Febbraio MA, Cameron-Smith D: 17beta-estradiol upregulates the expression of peroxisome proliferator-activated receptor and lipid oxidative genes in skeletal muscle. J Mol Endocrinol 2003.

72. Gao H, Bryzgalova G, Hedman E, Khan A, Efendic S, Gustafsson JA, Dahlman-Wright K: Long-term administration of estradiol decreases expression of hepatic lipogenic genes and improves insulin sensitivity in ob/ob mice: a possible mechanism is through direct regulation of signal transducer and activator of transcription 3. Mol Endocrinol 2006, 20(6):1287–1299.

73. Benedek G, Zhang J, Nguyen H, Kent G, Seifert HA, Davin S, Stauffer P, Vandenbark AA, Karstens L, Asquith M et al: Estrogen protection against EAE modulates the microbiota and mucosal-associated regulatory cells. J Neuroimmunol 2017, 310:51–59.

74. Lozupone CA, Stombaugh JI, Gordon JI, Jansson JK, Knight R: Diversity, stability and resilience of the human gut microbiota. Nature 2012, 489(7415):220–230.

75. Peters BA, Shapiro JA, Church TR, Miller G, Trinh-Shevrin C, Yuen E, Friedlander C, Hayes RB, Ahn J: A taxonomic signature of obesity in a large study of American adults. Sci Rep 2018, 8(1):9749.

76. Ormerod KL, Wood DL, Lachner N, Gellatly SL, Daly JN, Parsons JD, Dal’Molin CG, Palfreyman RW, Nielsen LK, Cooper MA et al: Genomic characterization of the uncultured Bacteroidales family S24-7 inhabiting the guts of homeothermic animals. Microbiome 2016, 4(1):36.

77. Chassard C, Bernalier-Donadille A: H2 and acetate transfers during xylan fermentation between a butyrate-producing xylanolytic species and hydrogenotrophic microorganisms from the human gut. FEMS Microbiol Lett 2006, 254(1):116–122.

78. Ríos-Covián D, Ruas-Madiedo P, Margolles A, Gueimonde M, de los Reyes-Gavilán CG, Salazar N: Intestinal Short Chain Fatty Acids and their Link with Diet and Human Health. Frontiers in Microbiology 2016, 7.

79. Macfarlane GT, Macfarlane S: Bacteria, colonic fermentation, and gastrointestinal health. Journal of AOAC International 2012, 95(1):50–60.

80. Krych L, Nielsen DS, Hansen AK, Hansen CH: Gut microbial markers are associated with diabetes onset, regulatory imbalance, and IFN-gamma level in NOD mice. Gut microbes 2015, 6(2):101–109.

81. Rooks MG, Veiga P, Wardwell-Scott LH, Tickle T, Segata N, Michaud M, Gallini CA, Beal C, van Hylckama-Vlieg JE, Ballal SA et al: Gut microbiome composition and function in experimental colitis during active disease and treatment-induced remission. ISME J 2014, 8(7):1403–1417.

82. Morgan XC, Tickle TL, Sokol H, Gevers D, Devaney KL, Ward DV, Reyes JA, Shah SA, LeLeiko N, Snapper SB et al: Dysfunction of the intestinal microbiome in inflammatory bowel disease and treatment. Genome Biol 2012, 13(9):R79.

83. Bäckhed F, Ding H, Wang T, Hooper LV, Koh GY, A. N, Semenkovich CF, Gordon JI: The gut microbiota as an environmental factor that regulates fat storage. Proc Natl Acad Sci U S A 2004, 101(44):15718–15723.

84. Hildebrandt MA, Hoffmann C, Sherrill-Mix SA, Keilbaugh SA, Hamady M, Chen YY, Knight R, Ahima RS, Bushman F, Wu GD: High-fat diet determines the composition of the murine gut microbiome independently of obesity. Gastroenterology 2009, 137(5):1716–1724 e1711-1712.

85. Zhang C, Zhang M, Pang X, Zhao Y, Wang L, Zhao L: Structural resilience of the gut microbiota in adult mice under high-fat dietary perturbations. ISME J 2012, 6(10):1848–1857.

86. Turnbaugh PJ, Ley RE, Mahowald MA, Magrini V, Mardis ER, Gordon JI: An obesity-associated gut microbiome with increased capacity for energy harvest. Nature 2006, 444(7122):1027–1031.

87. Semova I, Carten JD, Stombaugh J, Mackey LC, Knight R, Farber SA, Rawls JF: Microbiota regulate intestinal absorption and metabolism of fatty acids in the zebrafish. Cell Host Microbe 2012, 12(3):277–288.

88. Barengolts E, Green SJ, Eisenberg Y, Akbar A, Reddivari B, Layden BT, Dugas L, Chlipala G: Gut microbiota varies by opioid use, circulating leptin and oxytocin in African American men with diabetes and high burden of chronic disease. PLoS One 2018, 13(3):e0194171.

89. Christensen HR, Frokiaer H, Pestka JJ: Lactobacilli Differentially Modulate Expression of Cytokines and Maturation Surface Markers in Murine Dendritic Cells. J Immunol 2002, 168(1):171–178.

90. Madsen KL, Doyle JS, Jewell LD, Tavernini MM, Fedorak RN: Lactobacillus species prevents colitis in interleukin 10 gene-deficient mice. Gastroenterology 1999, 116(5):1107–1114.

91. Lepage P, Hasler R, Spehlmann ME, Rehman A, Zvirbliene A, Begun A, Ott S, Kupcinskas L, Dore J, Raedler A et al: Twin study indicates loss of interaction between microbiota and mucosa of patients with ulcerative colitis. Gastroenterology 2011, 141(1):227–236.

92. Martínez I, Perdicaro DJ, Brown AW, Susan Hammons S, Carden TJ, Carr TP, Eskridge KM, Walter J: Diet-Induced Alterations of Host Cholesterol Metabolism Are Likely To Affect the Gut Microbiota Composition in Hamsters. Appl Environ Microbiol 2013.

93. Zhang H, DiBaiseb JK, Zuccoloc A, Kudrnac D, Braidottic M, Yuc Y, Parameswarana P, Crowellb MD, Wingc R, Bruce E. Rittmanna BE et al: Human gut microbiota in obesity and after gastric bypass. Proc Natl Acad Sci U S A 2009, 106:2365–2370.

94. Jensen EV, Suzuki T, Kawashima T, Stumpf WE, Jungblut PW, DeSombre ER: A two-step mechanism for the interaction of estradiol with rat uterus. Proc Natl Acad Sci U S A 1968, 59(2):632–638.

95. Kuiper GG, Enmark E, Pelto-Huikko M, Nilsson S, Gustafsson JA: Cloning of a novel receptor expressed in rat prostate and ovary. Proc Natl Acad Sci U S A 1996, 93(12):5925–5930.

96. Tetel MJ, Pfaff DW: Contributions of estrogen receptor-alpha and estrogen receptor-ss to the regulation of behavior. Biochim Biophys Acta 2010, 1800(10):1084–1089.

97. Mani SK, Oyola MG: Progesterone signaling mechanisms in brain and behavior. Front Endocrinol (Lausanne) 2012, 3:7.

98. Frank A, Brown LM, Clegg DJ: The role of hypothalamic estrogen receptors in metabolic regulation. Front Neuroendocrinol 2014, 35(4):550–557.

99. Qiu J, Bosch MA, Tobias SC, Krust A, Graham SM, Murphy SJ, Korach KS, Chambon P, Scanlan TS, Ronnekleiv OK et al: A G-protein-coupled estrogen receptor is involved in hypothalamic control of energy homeostasis. J Neurosci 2006, 26(21):5649–5655.

100. Park CJ, Zhao Z, Glidewell-Kenney C, Lazic M, Chambon P, Krust A, Weiss J, Clegg DJ, Dunaif A, Jameson JL et al: Genetic rescue of nonclassical ERalpha signaling normalizes energy balance in obese Eralpha-null mutant mice. J Clin Invest 2011, 121(2):604–612.

101. Santollo J, Katzenellenbogen BS, Katzenellenbogen JA, Eckel LA: Activation of ERalpha is necessary for estradiol’s anorexigenic effect in female rats. Horm Behav 2010, 58(5):872–877.

102. Santollo J, Wiley MD, Eckel LA: Acute activation of ER alpha decreases food intake, meal size, and body weight in ovariectomized rats. Am J Physiol Regul Integr Comp Physiol 2007, 293(6):R2194–2201.

103. Heine PA, Taylor JA, Iwamoto GA, Lubahn DB, Cooke PS: Increased adipose tissue in male and female estrogen receptor-alpha knockout mice. Proc Natl Acad Sci U S A 2000, 97(23):12729–12734.

104. Menon R, Watson SE, Thomas LN, Allred CD, Dabney A, Azcarate-Peril MA, Sturino JM: Diet complexity and estrogen receptor beta status affect the composition of the murine intestinal microbiota. Appl Environ Microbiol 2013, 79(18):5763–5773.

105. Enmark E, Pelto-Huikko M, Grandien K, Lagercrantz S, Lagercrantz J, Fried G, Nordenskjold M, Gustafsson JA: Human estrogen receptor beta-gene structure, chromosomal localization, and expression pattern. J Clin Endocrinol Metab 1997, 82(12):4258–4265.

106. Campbell-Thompson M, Lynch IJ, Bhardwaj B: Expression of estrogen receptor (ER) subtypes and ERbeta isoforms in colon cancer. Cancer Res 2011.

107. McIntosh FM, Maison N, Holtrop G, Young P, Stevens VJ, Ince J, Johnstone AM, Lobley GE, Flint HJ, Louis P: Phylogenetic distribution of genes encoding beta-glucuronidase activity in human colonic bacteria and the impact of diet on faecal glycosidase activities. Environ Microbiol 2012, 14(8):1876–1887.

108. Kwa M, Plottel CS, Blaser MJ, Adams S: The Intestinal Microbiome and Estrogen Receptor-Positive Female Breast Cancer. J Natl Cancer Inst 2016, 108(8).

109. Flores R, Shi J, Fuhrman B, Xu X, Veenstra TD, Gail MH, Gajer P, Ravel J, Goedert JJ: Fecal microbial determinants of fecal and systemic estrogens and estrogen metabolites: a cross-sectional study. J Transl Med 2012, 10:253.

110. Cani PD, Bibiloni R, Knauf C, Waget A, Neyrinck AM, Delzenne NM, Burcelin R: Changes in gut microbiota control metabolic endotoxemia-induced inflammation in high-fat diet-induced obesity and diabetes in mice. Diabetes 2008, 57(6):1470–1481.

111. Cani PD, Amar J, Iglesias MA, Poggi M, Knauf C, Bastelica D, Neyrinck AM, Fava F, Tuohy KM, Chabo C et al: Metabolic endotoxemia initiates obesity and insulin resistance. Diabetes 2007, 56(7):1761–1772.

112. Blasco-Baque V, Serino M, Vergnes JN, Riant E, Loubieres P, Arnal JF, Gourdy P, Sixou M, Burcelin R, Kemoun P: High-fat diet induces periodontitis in mice through lipopolysaccharides (LPS) receptor signaling: protective action of estrogens. PLoS One 2012, 7(11):e48220.

113. Rajala MW, Patterson CM, Opp JS, Foltin SK, Young VB, Myers MG, Jr.: Leptin acts independently of food intake to modulate gut microbial composition in male mice. Endocrinology 2014, 155(3):748–757.

114. Ng KY, Yong J, Chakraborty TR: Estrous cycle in ob/ob and ovariectomized female mice and its relation with estrogen and leptin. Physiol Behav 2010, 99(1):125–130.

